# Polygenic Risk Scores for Cardio-renal-metabolic Diseases in the Penn Medicine Biobank

**DOI:** 10.1101/759381

**Authors:** R.L. Kember, A. Verma, S. Verma, A. Lucas, R. Judy, J. Chen, Regeneron Genetics Center, S. Damrauer, D.J. Rader, M.D. Ritchie

## Abstract

Cardio-renal-metabolic (CaReMe) conditions are common and the leading cause of mortality around the world. Genome-wide association studies have shown that these diseases are polygenic and share many genetic risk factors. Identifying individuals at high genetic risk will allow us to target prevention and treatment strategies. Polygenic risk scores (PRS) are aggregate weighted counts that can demonstrate an individual’s genetic liability for disease. However, current PRS are often based on European ancestry individuals, limiting the implementation of precision medicine efforts in diverse populations. In this study, we develop PRS for six diseases and traits related to cardio-renal-metabolic disease in the Penn Medicine Biobank. We investigate their performance in both European and African ancestry individuals, and identify genetic and phenotypic overlap within these conditions. We find that genetic risk is associated with the primary phenotype in both ancestries, but this does not translate into a model of predictive value in African ancestry individuals. We conclude that future research should prioritize genetic studies in diverse ancestries in order to address this disparity.

## Introduction

In this era of precision medicine, there are significant efforts to identify the genetic, environmental, family history, and clinical factors that influence the risk of disease as well as the influence of these factors on disease prognosis and treatment. Knowing in advance the factors that can lead to increased risk of disease can provide a major health benefit to individuals, as treatment and support strategies can be targeted towards individuals at higher risk. Identification of a large number of loci with small genetic effects in genome-wide association studies (GWAS) have highlighted the polygenic behavior of most common, complex diseases^1,2^. An emerging technology in the field of disease risk prediction is the polygenic risk score (PRS). PRS is the cumulative, mathematical aggregation of risk derived from the contributions of many DNA variants across the genome^3^.

Recent studies show high prevalence of cardio-renal-metabolic conditions among adults in the USA^4^ and together they are the leading cause of mortality around the world^5,6^. GWAS have identified more than 100 loci associated with common diseases such as coronary artery disease (CAD), body mass index (BMI), hypertension, renal failure and type 2 diabetes (T2D). This group of cardio, renal, and metabolic conditions are collectively referred to as CaReMe conditions. Among the individuals that are diagnosed with one disease (for example T2D), the prevalence of comorbidities such as hypertension, CAD, heart failure (HF), and chronic kidney disease (CKD) also increases. To evaluate disease risk in an individual, it is essential to also consider comorbid or secondary conditions related to the primary disease. There are several GWA-studies that have identified shared genetic associations between CaReMe conditions, demonstrating similarity in the underlying genetic architecture^7,8^. Pathophysiology of these conditions also show the cross-talk between organ systems and its effect on disease progression such as hemodynamic interaction between heart and kidney in heart failure^9^. With PRS, we can derive individuals’ disease risk for each CaReMe condition using GWAS summary statistics. More importantly, PRS is derived from the effect of millions of genetic variants on a disease; so it accounts for an individual’s genetic background. Therefore, PRS can evaluate the genetic overlap among coexisting or comorbid conditions. Phenome-wide Association Studies (PheWAS) can be used to identify links between disease risk and other conditions^10–12^. Using these strategies, we investigated whether cross-phenotype associations can provide insights into the contribution of risk for one disease risk on other conditions. Lastly, we also evaluated the effect of age, sex, and ancestry on CaReMe PRS predictions.

There are several strategies to derive PRS for a disease of interest. Traditionally, genetic risk scores (GRS) were derived using the genome-wide significant SNPs from a genome-wide association study; however, recent studies show that using association results with much lower p-value significance (p<0.05) segregate individuals risk with better accuracy^1^. The development and clinical utility of PRS is under active investigation, especially in racial and ethnic minority populations^13–15^. Most large-scale GWAS have been conducted in individuals from European descent populations and most PRS are derived from these studies. Subsequently, the majority of PRS investigations published to date have been conducted in populations of European ancestry^16^. There can be several differences such as linkage disequilibrium (LD) structure and allele frequency of the variants, which can lead to inaccurate PRS for non-European populations^16^. This is not unique to PRS studies, but the majority of human genetic research suffers from this same phenomenon^17^. In this study, we investigated the implementation of PRS for cardio-renal-metabolic conditions in European (EUR) and African (AFR) ancestry individuals in the Penn Medicine Biobank (PMBB). PMBB is a cohort of 52,853 individuals established for genomic and precision medicine research. Approximately 20,000 of the individuals in the study have genetic data from a genotyping array which has been imputed to the 1000Genomes phase III using the Michigan Imputation Server^18^. Approximately 25% of the PMBB study population is African ancestry. We calculated PRS in the PMBB genetic data to evaluate 1) risk prediction accuracy among EUR and AFR based on GWAS summary statistics generated in EUR data; and 2) the utility of PRS in determining genetic overlap among CaReMe conditions.

## Methods

### Penn Medicine Biobank

The Penn Medicine BioBank (PMBB) recruits participants through the University of Pennsylvania Health System by enrolling at the time of appointment. Patients participate by donating either blood or a tissue sample and allowing researchers access to their electronic health record (EHR) information. This academic biobank provides researchers with centralized access to a large number of blood and tissue samples with attached health information. The facility banks both blood specimens (i.e., whole blood, plasma, serum, buffy coat, and DNA isolated from leukocytes) and tissues (i.e., formalin-fixed paraffin-embedded, fresh and flash frozen). PMBB currently consists of 52,853 consented samples. Approximately one third (N=19,515) of these participants have been genotyped to date. PMBB is a diverse cohort, with 70% European ancestry, 25% African ancestry, and 5% Asian or Latino ancestry. See Table 1 for characteristics of all participants.

**Table 1.** Participant Characteristics

|  | **PMBB consented patients** | **PMBB genotyped patients** |
| --- | --- | --- |
| Total Patients | 52,853 | 19,515 |
| Female, (%) | 25,926 (49%) | 7856 (41%) |
| Age (average) | 18 – 99 (60) | 20 – 99 (66) |
| Body mass index | 29.36 (13 - 83) | 30.02 (13 - 83) |
| **Race** |  |  |
| American Indian or Alaska Native | 34 | 1 |
| Asian | 1,201 | 158 |
| Black or African American | 11,173 | 6159 |
| Native Hawaiian or other Pacific Islander | 30 | 7 |
| Other | 1,577 | 459 |
| Unknown | 1,775 | 533 |
| White | 36,707 | 10,563 |
| **Ethnicity** |  |  |
| Hispanic or Latino | 1,381 | 350 |
| Not Hispanic or Latino | 50,994 | 17,517 |
| Unknown | 122 | 13 |

### Genotyping and Quality Control and Imputation

DNA extracted from the blood plasma of 19,515 samples were genotyped in three batches: 10,867 samples on the Illumina QuadOmni chip at the Regeneron Genetics Center; 5,676 samples on the Illumina GSA V1 chip and 2,972 samples on the Illumina GSA V2 chip by the Center for Applied Genomics at the Children’s Hospital of Philadelphia. Due to the low overlap among genetic variants on the different genotyping arrays, we used an imputation strategy to combine these datasets^18,19^. Prior to imputation, we applied a quality control pipeline^19^ to each dataset, removing individuals with sex errors or had a sample call rate <90%; and removing variants which were palindromic or had a call rate <95%. Table 2 summarizes each dataset before and after QC.

**Table 2:** Summary of genotype data before and after QC.

| Dataset | Pre-QC |  | Post-QC |  |
| --- | --- | --- | --- | --- |
|  | #Samples | #SNPs | #Samples | #SNPs |
| Illumina Infinium OMNI | 10,867 | 713,599 | 10,506 | 651,366 |
| Illumina GSA V1 | 5,676 | 700,078 | 5,660 | 666,032 |
| Illumina GSA V2 | 2,972 | 759,993 | 2,965 | 700,984 |

Genotypes for each of the three PMBB datasets were phased (Eagle v2.3) and imputed to the 1000Genomes reference panel (1000G Phase3 v5) using the Michigan Imputation Server^18^. Accuracy of the imputed variants was assessed via comparison of the expected vs actual allele frequency of variants (R^2^=0.3). Following imputation, the datasets were merged, with each position matched based on alleles. In the merged dataset, the average R^2^ of variants = 0.75. Genetic ancestry was calculated from common, high-quality SNPs (MAF > 0.05, missingness < 0.1) using SMARTPCA^20^ module of the Eigensoft package. We split the merged file into individuals with European ancestry (N=11,524) and individuals with African ancestry (N=5,994). All subsequent QC and analysis steps were performed independently within each population.

We retained high quality, common SNPs with imputation marker R2 ≥ 0.7 and minor allele frequencies ≥ 0.01. We identified and removed related individuals using a kinship coefficient of 0.25. Using a graph-based algorithm, we selected and removed the sample that is closely related to the most samples within the set of related samples. Following QC, we retained 10,351 European ancestry individuals and 5,553 African ancestry individuals. Ancestry specific principal components were generated within each ancestral group following ancestry assignment, and these were used as covariates for subsequent analyses. Genetic ancestry of individuals was determined by performing quantitative discriminate analyses on PCs.

### Polygenic Risk Scores

To derive PRS, we used the summary statistics from the largest and/or most recent GWAS studies for each trait (See Table 3). To reduce our total SNP set to a size amenable for PRS analysis, we first extracted SNPs present in the HapMap reference panel (N SNPs =1,437,731 in HapMap panel; retained 1,320,405 SNPs in EUR and AFR datasets). LDPred (v1.0) was used to generate the posterior mean effect of each SNP based on the LD information from either the PMBB European dataset or the PMBB African dataset^21^. PRS on each population were calculated using PLINK v1.9^22^. We tested several values for LDPred’s tuning parameter “fraction of causal variants” (*ρ*=0.001, 0.003, 0.01, 0.03, 0.1, 0.3, 1) for deriving SNP weights.

**Table 3:** Source of GWAS used for PRS creation

| **Disease** | **GWAS source** | **Sample size in GWAS** | **Reference** |
| --- | --- | --- | --- |
| Type 2 Diabetes | DIAGRAM | 26,488 cases | PMID: 24509480 |
| Body Mass Index | GIANT | 339,224 | PMID: 25673413 |
| Hypertension | UK Biobank | 144,793 | PMID: 30940143 |
| Myocardial Infarction | CARDIoGRAMplus C4D consortium | 43,154 cases | PMID: 26343387 |
| Coronary Artery Disease | CARDIoGRAMplus C4D consortium | 60,801 cases | PMID: 26343387 |
| Chronic Kidney Disease | CKD Genetics consortium | 41,395 cases | PMID: 31152163 |

### Phenotypes

We derived phenotypes using ICD-9 and ICD-10 data for 52,853 individuals from the electronic health record (EHR), consisting of 11.8 million records. We filtered on encounter type to identify records representing encounters with a physician (see Supplemental Table 1 for encounters selected). ICD-9 codes were aggregated to phecodes using the phecode ICD9 map 1.2^10,23^; ICD-10 codes were aggregated to phecodes using the phecode ICD-10cm map 1.2 (beta)^24^. Individuals are considered cases for the phenotype if they have at least 2 instances of the phecode on unique dates, controls if they have no instance of the phecode, and ‘other/missing’ if they have one instance or a related phecode. By the following criteria, there were a total of 1,812 phecodes included in the analysis.

### Statistical Analysis

PRS were standardized with mean = 0 and SD = 1. Logistic regression models accounting for age, sex, and the first 10 within-ancestry principal components (PCs) were used to test for association of PRS with the primary phenotype. Area under the receiver operator curve (AUC) was determined using the R package pROC, using the same logistic regression model as above. AUC was also calculated for covariates alone.

A Phenome-wide Association Study (PheWAS) was performed for the optimal PRS identified in the above analysis for each primary condition. Logistic regression models with each PRS as the independent variable, phecodes as the dependent variables, and age, sex, and the first 10 PCs as covariates were used to identify secondary phenotypic associations. A phenome-wide bonferroni significance threshold of 2.7 × 10^−5^ was applied to account for multiple testing.

## Results

### Demographics of PMBB dataset

Using phecodes, we identified 7,476 EUR ancestry individuals (73.4%) and 4,177 AFR ancestry individuals (76.4%) with either type 2 diabetes, obesity, hypertension, myocardial infarction, coronary atherosclerosis, or renal failure (Table 4). In EUR ancestry individuals, 24.7% had been diagnosed with one instance of disease, 35.2% had 2-3 diseases, 12.5% had 4-5 diseases, and 1.0% had all six diseases. In AFR ancestry individuals, 20.8% had been diagnosed with one instance of disease, 39.1% had 2-3 diseases, 14.6% had 4-5 diseases, and 1.9% had all six diseases.

**Table 4:** Counts for phecodes in the Penn Medicine Biobank

| **Phecode** | **Description** | **# EUR**<br>**(total=10,182)** | **# AFR (total=5,465)** |
| --- | --- | --- | --- |
| 250.2 | Type 2 diabetes | 2,119 | 1,804 |
| 278.1 | Obesity | 1,720 | 2,107 |
| 401 | Hypertension | 5,980 | 3,455 |
| 411.2 | Myocardial Infarction | 1,193 | 469 |
| 411.4 | Coronary atherosclerosis | 3,996 | 975 |
| 585 | Renal failure | 2,191 | 1,635 |

### Determining the PRS with the best discriminative capacity

We generated a PRS for each phenotype of interest: type 2 diabetes, body mass index, hypertension, myocardial infarction, coronary artery disease, and chronic kidney disease (see methods, Table 3). Candidate PRS were generated for 7 parameters, and their association with the primary phenotype tested. All PRS had at least one parameter that was significantly associated with their primary phenotype (Figure 1, Supplemental Table 2). We selected the best performing PRS based on the maximum area under the receiver operator curve (AUC; Supplemental Table 3). Type 2 diabetes PRS was significantly associated with type 2 diabetes (best parameter for EUR *ρ*=0.01, OR=1.52, p=6.62×10^−43^, best parameter for AFR *ρ*=0.01, OR=1.3, p=2.19×10^−13^). BMI PRS was significantly associated with obesity (best parameter for EUR *ρ*=0.3, OR=1.7, p=8.97×10^−65^, best parameter for AFR *ρ*=0.1, OR=1.2, p=5.55×10^−10^). Hypertension PRS was significantly associated with hypertension (best parameter for EUR *ρ*=1, OR=1.4, p=4.50×10^−40^, best parameter for AFR *ρ*=0.3, OR=1.27, p=4.31×10^−10^). Myocardial infarction PRS was significantly associated with myocardial infarction (best parameter for EUR *ρ*=0.01, OR=1.8, p=3.74×10^−51^, best parameter for AFR *ρ*=0.1, OR=1.3, p=5.01×10^−7^). Coronary artery disease PRS was significantly associated with coronary atherosclerosis (best parameter for EUR *ρ*=0.01, OR=1.66, p=1.54×10^−77^, best parameter for AFR *ρ*=0.03, OR=1.27, p=2.53×10^−8^). Chronic kidney disease PRS was significantly associated with renal failure (best parameter for EUR *ρ*=0.01, OR=1.2, p=2.60×10^−6^, best parameter for AFR *ρ*=0.001, OR=1.1, p=0.024).

**Figure 1.**
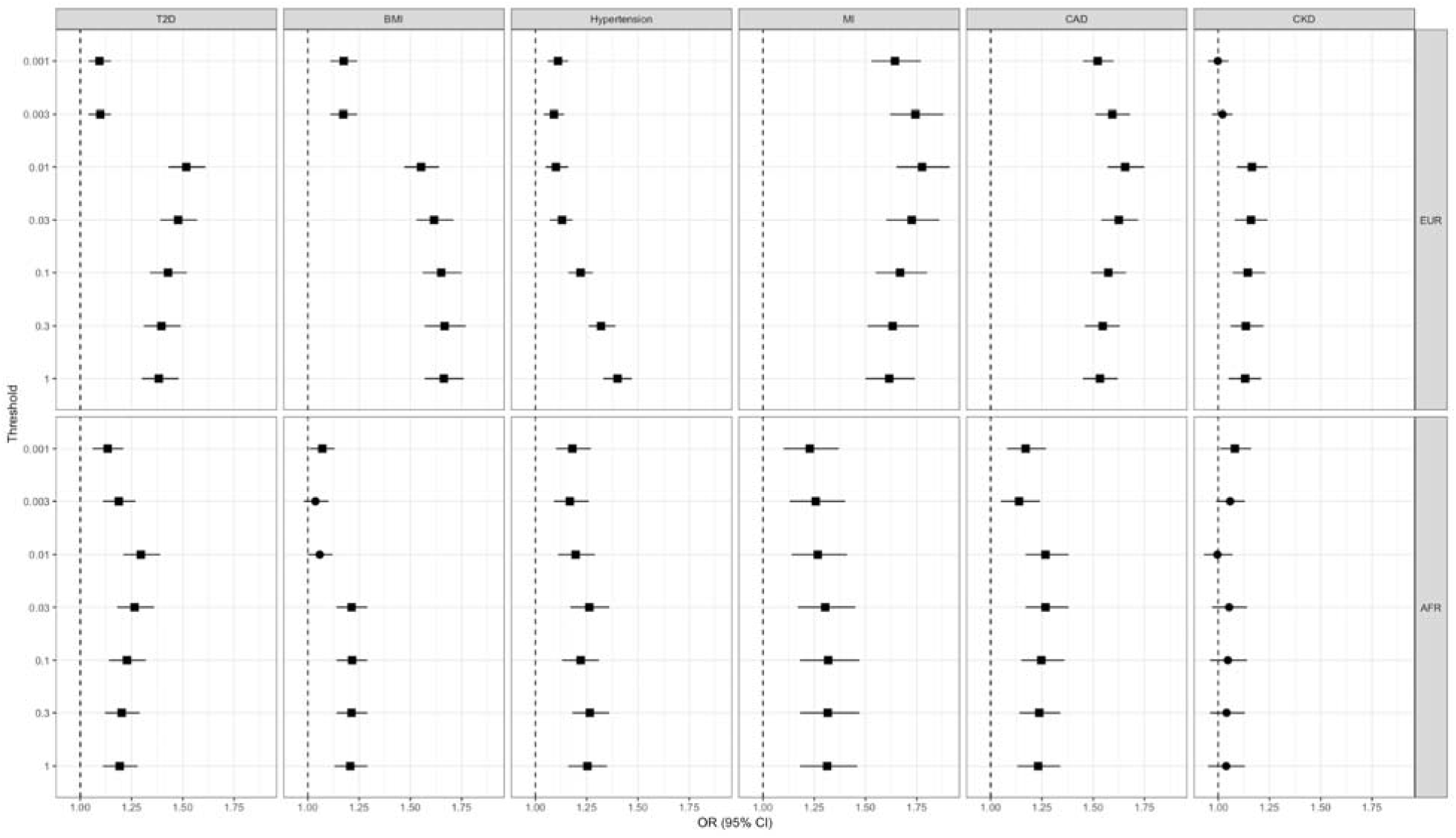
Association of PRS with primary phenotype for EUR and AFR ancestry. Square denotes p<0.05.

### Performance of PRS in EUR vs. AFR ancestry

We selected the parameter that produced the strongest associated candidate PRS for each PRS-phenotype grouping within each ancestry for further analysis. In all cases, PRS performance was best in European ancestry individuals. The distribution of PRS in cases and controls in both populations is illustrated in Figure 2. In European ancestry, the mean distribution of PRS in cases is consistently higher than controls. In African ancestry individuals this difference is much smaller, with substantial overlap between the PRS distribution in cases and controls. This is reflected in the comparison between the AUC for the full model and the AUC for covariates alone (Supplemental Table 3). Although the AUC in the full model is high in both ancestries (0.57-0.84), showing ability to distinguish between cases and controls, in AFR the full model offers little improvement over the model based on covariates alone (average improvement in AUC for best PRS=0.007). In contrast, in EUR the covariate model is improved when the PRS is added (average improvement in AUC for best PRS=0.032). Further, to evaluate the significance of variables in the full model (PRS and covariates), we performed a step-wise regression. In AFR, we identified that PRS was not selected in the best model in CAD but was selected in other phenotypes. CAD step-wide regression model further support our argument that PRS derived from EUR GWAS studies might not add any risk prediction in AFR individuals (Supplementary Table 4).

**Figure 2.**
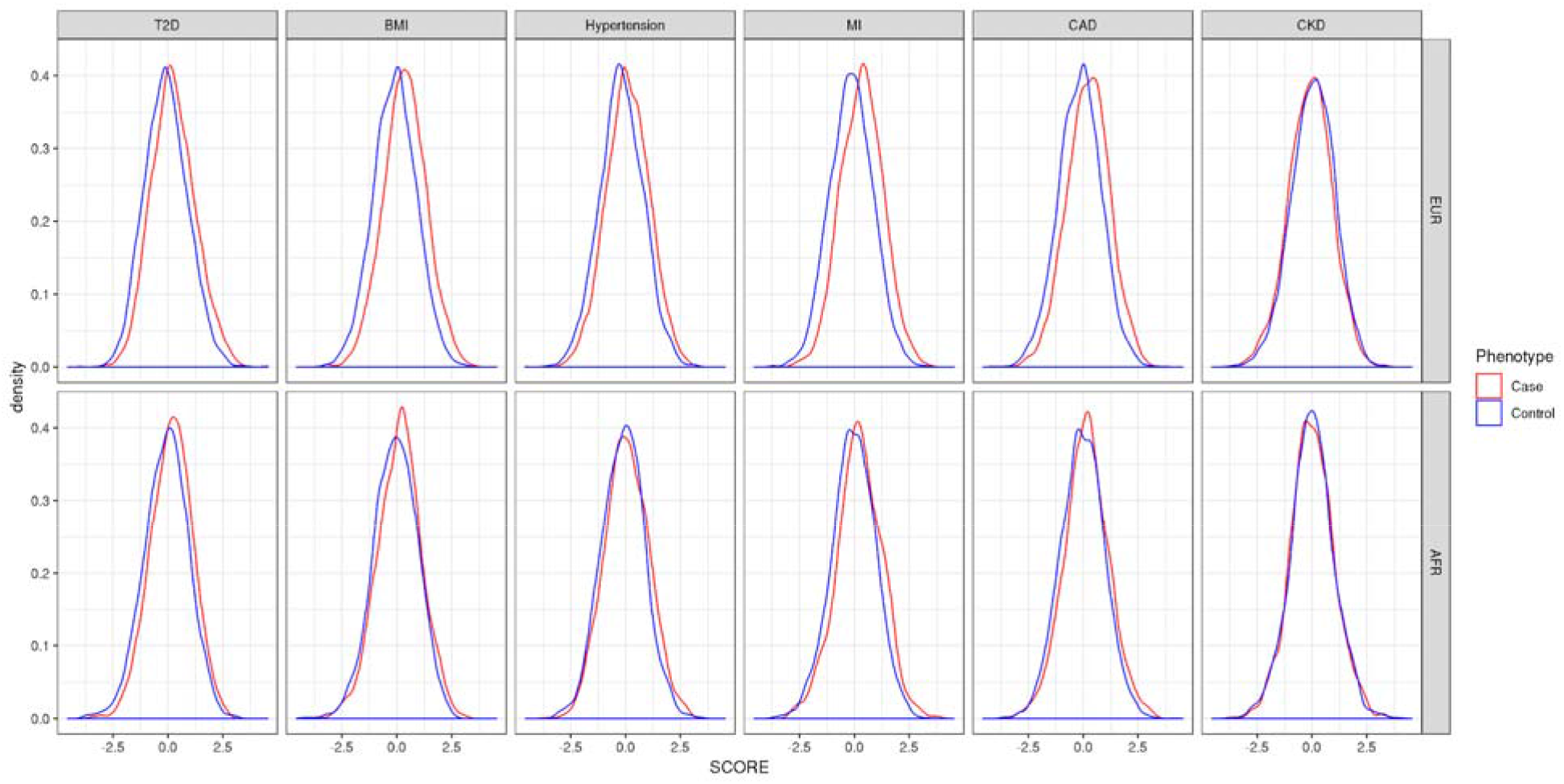
Density plot of polygenic risk scores in cases (red) and controls (blue) in both European and African ancestry individuals.

### Correlation of PRS

We next tested for correlations between the polygenic risk scores for the six selected diseases in each ancestry (Figure 3, Supplemental Table 5). In European ancestry individuals, we identified significant positive correlations between CKD/T2D, CKD/BMI, CAD/BMI, CAD/Hypertension, CAD/MI, MI/BMI, MI/Hypertension, Hypertension/T2D, Hypertension/BMI, and BMI/T2D. CKD PRS was also negatively correlated with PRS for MI and CAD. In African ancestry individuals, we identified significant positive correlations between CKD/BMI, CAD/T2D, CAD/BMI, CAD/Hypertension, CAD/MI, MI/T2D, MI/BMI, MI/Hypertension, Hypertension/T2D, Hypertension/BMI, and BMI/T2D. Overall, the correlations identified in PRS in African ancestry individuals were weaker than those identified in PRS in European ancestry individuals.

**Figure 3.**
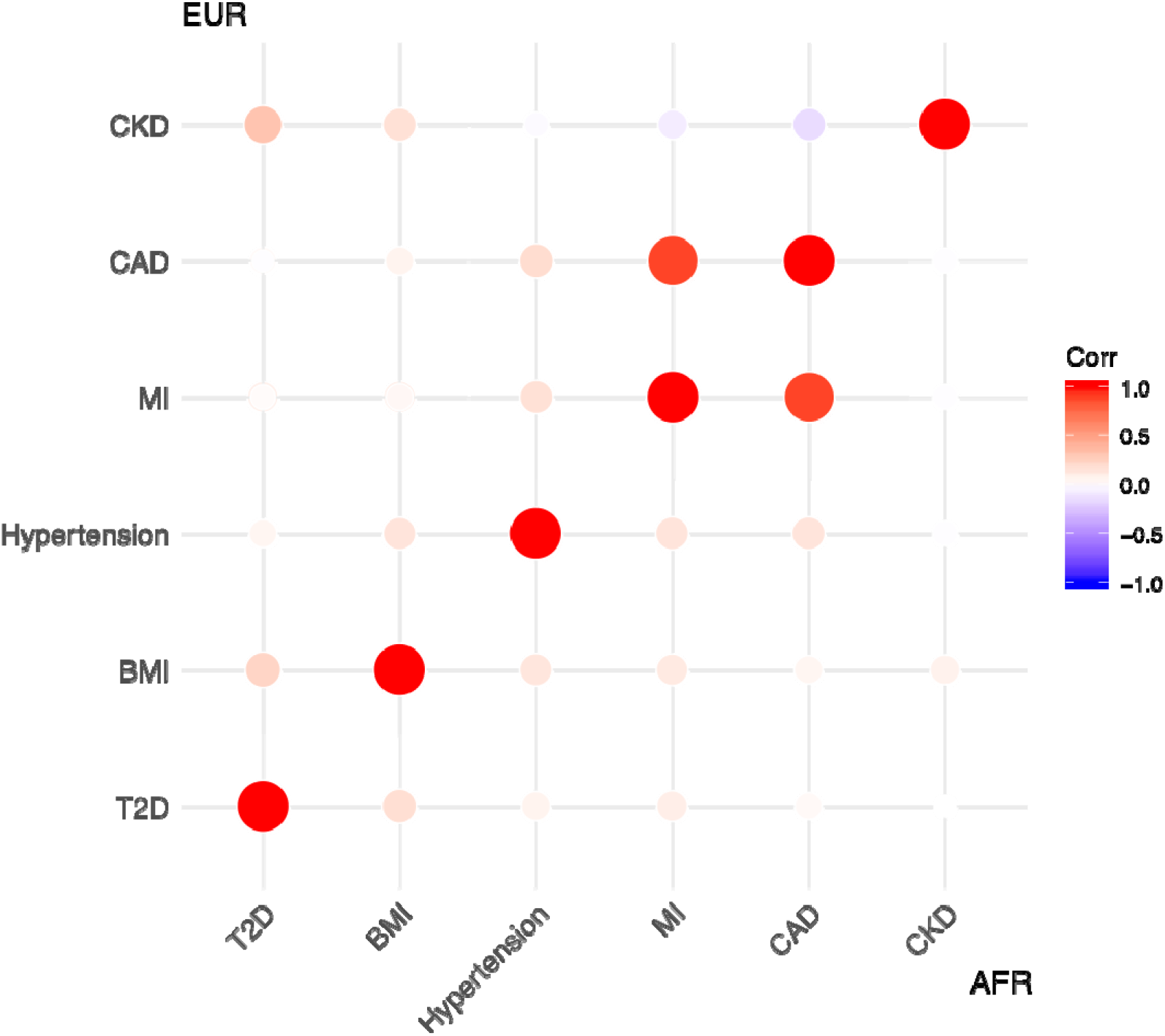
Correlation between polygenic risk scores for six diseases in European and African ancestry individuals. The size of the circle denotes the size of the correlation. The color denotes the direction of correlation (red=positive, blue=negative). White circles denote a non-significant association. Correlation of European PRS are shown in the upper left, correlation of African PRS are shown in the lower right.

### Association of PRS with disease burden in cases

Using linear regression in cases only, we tested whether increased PRS is associated with increased burden of cardio-renal-metabolic disease. All PRS, except for CKD PRS, were significantly associated with increased disease burden in both European and African ancestry individuals (Table 5). In Europeans, BMI PRS was the strongest association with occurrence of multiple diseases (beta=0.14, p=6.6×10^−19^), in contrast to in African ancestry individuals, where the association remained significant but was reduced in effect size (beta=0.05, p=0.02). In African ancestry individuals, MI PRS was the strongest association with increased disease burden (beta=0.09, p=8.3×10^−7^).

**Table 5.** Association of PRS with the burden of disease in European and African ancestry individuals.

| PRS | EUR |  |  | AFR |  |  |
| --- | --- | --- | --- | --- | --- | --- |
|  | Beta | SE | P | Beta | SE | P |
| T2D | 0.117 | 0.017 | $5.47 \times 10^{-12}$ | 0.090 | 0.020 | $7.18 \times 10^{-6}$ |
| BMI | 0.135 | 0.015 | $6.56 \times 10^{-19}$ | 0.046 | 0.020 | 0.020 |
| Hypertension | 0.112 | 0.015 | $2.40 \times 10^{-14}$ | 0.037 | 0.019 | 0.048 |
| MI | 0.115 | 0.015 | $3.85 \times 10^{-15}$ | 0.093 | 0.019 | $8.28 \times 10^{-7}$ |
| CAD | 0.131 | 0.015 | $6.31 \times 10^{-18}$ | 0.092 | 0.019 | $1.10 \times 10^{-6}$ |
| CKD | 0.027 | 0.019 | 0.146 | 0.020 | 0.019 | 0.300 |

### PheWAS of polygenic risk scores reveals secondary associations

We performed a PheWAS of each PRS in both European and African ancestry individuals to identify secondary phenotypes associated with genetic risk (Figure 4, Supplemental Tables 6-11). In all PRS, associations with secondary phenotypes in AFR individuals were reduced. All PRS except CKD PRS were associated with secondary phenotypes. T2D PRS was associated with hypertension (OR=1.13, p=9.4×10^−6^) in EUR, and with Type 1 diabetes (OR=1.45, p=4.8×10^−6^) in AFR. BMI PRS was associated with multiple circulatory system phenotypes in EUR, including hypertension (OR=1.17, p=1.4×10^−9^), heart failure (OR=1.13, p=1.1×10^−6^), and coronary atherosclerosis (OR=1.12, p=5.3×10^−6^). BMI PRS was also associated with type 2 diabetes (OR=1.23, p=1.2×10^−14^), renal failure (OR=1.13, p=3.1×10^−6^), osteoporosis (OR=0.81, p=5.3×10^−5^), and sleep apnea (OR=1.26, p=4.3×10^−15^) in EUR. In AFR, the only additional phenotypes associated with BMI PRS were sleep apnea (OR=1.20, p=1.1×10^−6^) and use of an insulin pump (OR=1.28, p=8.6×10^−6^). Hypertension PRS was associated with circulatory system phenotypes in EUR, as well as type 2 diabetes (OR=1.29, p=6.8×10^−23^), disorders of lipoid metabolism (OR=1.16, p=2.6×10^−10^), renal failure (OR=1.13, p=3.2×10^−6^), and sleep apnea (OR=1.15, p=3.7×10^−7^). In AFR, hypertension PRS was not significantly associated with additional phenotypes. MI PRS was associated with hyperlipidemia (OR=1.22, p=1.8×10^−16^), hypertensive chronic kidney disease (OR=1.26, p=5.2×10^−9^) and type 2 diabetes (OR=1.12, p=9.4×10^−6^) in EUR, but no additional phenotypes in AFR. CAD PRS is associated with hyperlipidemia (OR=1.29, p=3.1×10^−24^), hypertensive chronic kidney disease (OR=1.28, p=1.4×10^−9^), hypertension (OR=1.16, p=6.3×10^−9^), type 2 diabetes (OR=1.14, p=9.6×10^−7^), and disorders of eye (OR=0.77, p=8.1×10^−5^) in EUR. In AFR, CAD PRS is associated with eye inflammation (OR=0.76, p=6.6×10^−5^).

**Figure 4.**
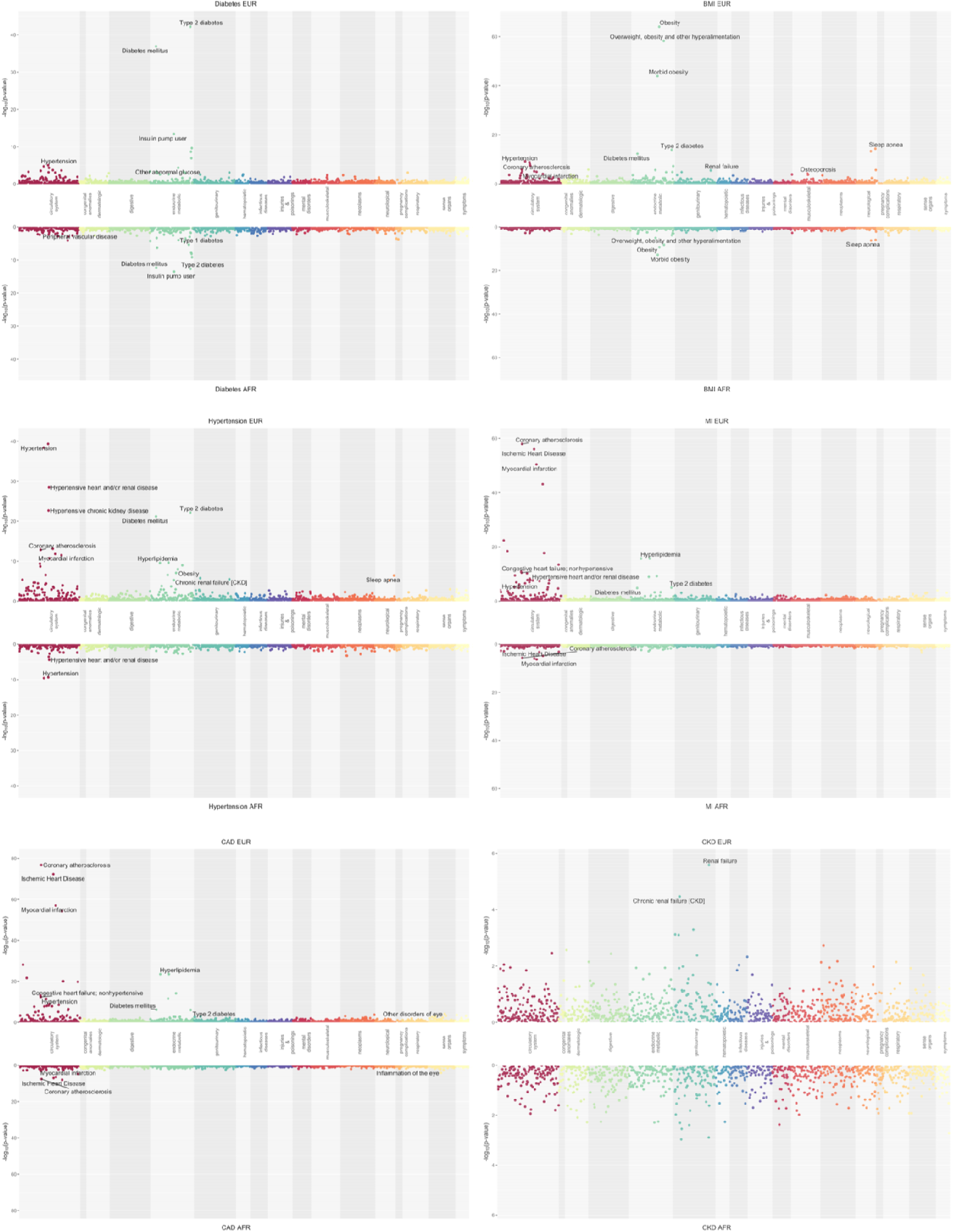
Secondary associations of polygenic risk for T2D, BMI, Hypertension, MI, CAD and CKD in EUR (top panels) and AFR (bottom panels) individuals.

## Discussion

We generated six polygenic risk scores representing genetic liability for cardio-renal-metabolic diseases, and investigated their performance in both European and African ancestry individuals in the Penn Medicine BioBank (PMBB), a biobank linked with electronic health records. For all phenotypes tested, we identified a significant association between the PRS and the primary phenotype in both ancestry groups. However, the ability of the PRS to discriminate between cases and controls varied among phenotypes. Furthermore, none of the PRS in AFR were sufficient to stratify individuals according to risk.

In European ancestry individuals, the PRS with the largest effect size was myocardial infarction, followed by coronary artery disease. The two GWAS that were used to generate these PRS also had the largest number of cases, and the PMBB dataset also contained a large number of cases for both of these diseases. However, the CKD PRS was the weakest performer in terms of effect size, despite it being based on the next largest GWAS and the PMBB containing a large number of individuals with renal failure. Therefore, while case number in both the GWAS and the target sample are clearly important, we believe that other factors such as disease heterogeneity, prevalence, penetrance, and non-additive effects among others must also play a role in the ability of PRS to associate with disease.

We conducted a number of analyses to explore secondary phenotypes associated with each PRS. First, we show that the PRS generated for the six diseases are correlated with each other, a finding supported by prior studies showing genetic correlations between CaReMe conditions^4^. We next show that increased PRS is associated with increased burden of disease, suggesting that a higher PRS burden may contribute in a non-discriminating fashion to disease outcome. Finally, we perform PheWAS analysis to identify secondary phenotypes associated with genetic liability for CaReMe diseases. Many of the secondary phenotypes identified could be attributed to the broader effects of disease risk factors and known comorbidities. For instance, risk for Type 2 diabetes was associated with hypertension, a known commonly co-occurring trait^25^. The BMI PRS was associated with sleep apnea, diabetes, hypertension and osteoporosis; all traits known to increase in individuals with higher BMI^26–29^. The extent to which these secondary phenotypes reflect causal associations between genetic risk and disease is unclear due to the commonality of co-morbidity of these traits.

Our findings highlight a major issue in the future implementation of PRS in clinical care. While GWAS conducted in EUR populations can be used to generate PRS that are associated with phenotype in AFR individuals, the scores generated are not sufficient to differentiate between cases and controls in a predictive model. This was an expected finding, and has been discussed widely in recent years as being a critical source of disparity in genetic research^14,16^. Due to differences in linkage disequilibrium patterns, effect sizes, and causal variants themselves, conducting GWAS in populations that are reflective of the patient population are necessary and will need to be prioritized in the coming years.

Finally, while there is much excitement and enthusiasm about PRS for clinical care, there is still significant research to be conducted to determine its optimal implementation. One of the most essential needs is to investigate how PRS can be incorporated alongside information commonly used to predict patients’ risk, such as family history, clinical comorbidities, and environmental/lifestyle factors. Many chronic diseases have published clinical guidelines with risk reduction recommendations (for example CVD^30^). The ultimate clinical utility of PRS will come to fruition when we understand how to integrate PRS with these published guidelines.

## Supporting information

Encounter Type

PheWAS of T2D PRS

PheWAS of BMI PRS

PheWAS of Hypertension PRS

PheWAS of MI PRS

PheWAS of CAD PRS

PheWAS of CKD PRS

## Supplemental Information

Supplemental Table 1: Encounter Type (see excel file)

**Supplemental Table 2:** Association of parameters of each PRS with their primary phenotype in EUR and AFR ancestry.

| Disease/PRS | Parameter | EUR |  | AFR |  |
| --- | --- | --- | --- | --- | --- |
|  |  | OR (95% CI) | P-value | OR (95% CI) | P-value |
| Type 2 diabetes/<br>Type 2 diabetes | $\rho=0.001$ | 1.09 (1.04-1.15) | $4.73 \times 10^{-4}$ | 1.13 (1.06-1.21) | $1.57 \times 10^{-4}$ |
| | $\rho=0.003$ | 1.1 (1.04-1.15) | $2.33 \times 10^{-4}$ | 1.19 (1.11-1.27) | $2.85 \times 10^{-7}$ |
| | $\rho=0.01$ | **1.52 (1.43-1.61)** | **$6.62 \times 10^{-43}$** | **1.3 (1.21-1.39)** | **$2.19 \times 10^{-13}$** |
| | $\rho=0.03$ | 1.48 (1.39-1.57) | $2.76 \times 10^{-34}$ | 1.26 (1.18-1.36) | $1.94 \times 10^{-10}$ |
| | $\rho=0.1$ | 1.43 (1.34-1.52) | $5.69 \times 10^{-27}$ | 1.23 (1.14-1.32) | $6.25 \times 10^{-8}$ |
| | $\rho=0.3$ | 1.4 (1.31-1.49) | $1.37 \times 10^{-23}$ | 1.2 (1.12-1.29) | $1.30 \times 10^{-6}$ |
| | $\rho=1$ | 1.38 (1.3-1.48) | $2.02 \times 10^{-22}$ | 1.19 (1.11-1.28) | $3.83 \times 10^{-6}$ |
| Obesity/<br>BMI | $\rho=0.001$ | 1.18 (1.11-1.24) | $2.81 \times 10^{-9}$ | 1.07 (1.01-1.13) | 0.022 |
| | $\rho=0.003$ | 1.17 (1.11-1.24) | $5.22 \times 10^{-9}$ | 1.04 (0.98-1.1) | 0.222 |
| | $\rho=0.01$ | 1.55 (1.47-1.64) | $6.85 \times 10^{-55}$ | 1.06 (1-1.12) | 0.059 |
| | $\rho=0.03$ | 1.62 (1.53-1.71) | $6.77 \times 10^{-62}$ | 1.21 (1.14-1.29) | $3.58 \times 10^{-10}$ |
| | $\rho=0.1$ | 1.65 (1.56-1.75) | $2.95 \times 10^{-64}$ | **1.22 (1.14-1.29)** | **$5.55 \times 10^{-10}$** |
| | $\rho=0.3$ | **1.67 (1.57-1.77)** | **$8.97 \times 10^{-65}$** | 1.21 (1.14-1.29) | $1.53 \times 10^{-9}$ |
| | $\rho=1$ | 1.66 (1.57-1.76) | $2.76 \times 10^{-63}$ | 1.21 (1.13-1.29) | $5.17 \times 10^{-9}$ |
| Hypertension/<br>Hypertension | $\rho=0.001$ | 1.11 (1.06-1.16) | $2.21 \times 10^{-5}$ | 1.18 (1.1-1.27) | $1.08 \times 10^{-5}$ |
| | $\rho=0.003$ | 1.09 (1.04-1.14) | $6.03 \times 10^{-4}$ | 1.17 (1.09-1.26) | $3.45 \times 10^{-5}$ |
| | $\rho=0.01$ | 1.10 (1.05-1.16) | $8.96 \times 10^{-5}$ | 1.20 (1.11-1.29) | $1.93 \times 10^{-6}$ |
| | $\rho=0.03$ | 1.13 (1.07-1.18) | $1.27 \times 10^{-6}$ | 1.26 (1.17-1.36) | $7.27 \times 10^{-10}$ |
| | $\rho=0.1$ | 1.22 (1.16-1.28) | $6.43 \times 10^{-16}$ | 1.22 (1.13-1.31) | $1.21 \times 10^{-7}$ |
| | $\rho=0.3$ | 1.32 (1.26-1.39) | $2.85 \times 10^{-29}$ | **1.27 (1.18-1.36)** | **$4.31 \times 10^{-10}$** |
| | $\rho=1$ | **1.40 (1.33-1.47)** | **$4.50 \times 10^{-40}$** | 1.25 (1.16-1.35) | $1.94 \times 10^{-9}$ |
| Myocardial<br>Infarction/<br>Myocardial<br>Infarction | $\rho=0.001$ | 1.64 (1.53-1.77) | $2.22 \times 10^{-41}$ | 1.23 (1.1-1.37) | $1.39 \times 10^{-4}$ |
| | $\rho=0.003$ | 1.74 (1.62-1.88) | $2.02 \times 10^{-49}$ | 1.26 (1.13-1.4) | $2.13 \times 10^{-5}$ |
| | $\rho=0.01$ | **1.78 (1.65-1.91)** | **$3.74 \times 10^{-51}$** | 1.27 (1.14-1.41) | $9.88 \times 10^{-6}$ |
| | $\rho=0.03$ | 1.73 (1.6-1.86) | $1.05 \times 10^{-45}$ | 1.3 (1.17-1.45) | $8.88 \times 10^{-7}$ |
| | $\rho=0.1$ | 1.67 (1.55-1.8) | $4.29 \times 10^{-40}$ | **1.32 (1.18-1.47)** | **$5.01 \times 10^{-7}$** |
| | $\rho=0.3$ | 1.63 (1.51-1.76) | $7.15 \times 10^{-37}$ | 1.32 (1.18-1.47) | $7.38 \times 10^{-7}$ |
| | $\rho=1$ | 1.62 (1.5-1.74) | $1.77 \times 10^{-35}$ | 1.31 (1.18-1.46) | $9.68 \times 10^{-7}$ |
| Coronary atherosclerosis/<br>coronary artery disease | $\rho=0.001$ | 1.52 (1.45-1.6) | $1.44 \times 10^{-60}$ | 1.17 (1.08-1.27) | $2.59 \times 10^{-4}$ |
| | $\rho=0.003$ | 1.59 (1.51-1.68) | $6.12 \times 10^{-71}$ | 1.14 (1.05-1.24) | 0.003 |
| | $\rho=0.01$ | **1.66 (1.57-1.75)** | **$1.54 \times 10^{-77}$** | 1.27 (1.17-1.38) | $2.87 \times 10^{-8}$ |
| | $\rho=0.03$ | 1.62 (1.54-1.72) | $4.53 \times 10^{-69}$ | **1.27 (1.17-1.38)** | **$2.53 \times 10^{-8}$** |
| | $\rho=0.1$ | 1.57 (1.49-1.66) | $8.72 \times 10^{-59}$ | 1.25 (1.15-1.36) | $1.89 \times 10^{-7}$ |
| | $\rho=0.3$ | 1.55 (1.46-1.63) | $6.01 \times 10^{-54}$ | 1.24 (1.14-1.34) | $5.25 \times 10^{-7}$ |
| | $\rho=1$ | 1.53 (1.45-1.62) | $1.04 \times 10^{-51}$ | 1.23 (1.13-1.34) | $9.05 \times 10^{-7}$ |
| Renal failure/<br>chronic kidney disease | $\rho=0.001$ | 1 (0.95-1.05) | 0.935 | **1.08 (1.01-1.16)** | **0.024** |
| | $\rho=0.003$ | 1.02 (0.97-1.07) | 0.408 | 1.06 (0.99-1.13) | 0.106 |
| | $\rho=0.01$ | **1.16 (1.09-1.24)** | **$2.60 \times 10^{-6}$** | 1 (0.93-1.07) | 0.908 |
| | $\rho=0.03$ | 1.16 (1.08-1.24) | $1.91 \times 10^{-5}$ | 1.05 (0.97-1.14) | 0.216 |
| | $\rho=0.1$ | 1.14 (1.07-1.23) | $1.48 \times 10^{-4}$ | 1.05 (0.96-1.14) | 0.285 |
| | $\rho=0.3$ | 1.13 (1.06-1.22) | $3.99 \times 10^{-4}$ | 1.04 (0.96-1.13) | 0.356 |
| | $\rho=1$ | 1.13 (1.05-1.21) | $5.61 \times 10^{-4}$ | 1.04 (0.95-1.13) | 0.379 |

**Supplemental Table 3:** AUC of parameters of each PRS with their primary phenotype in EUR and AFR ancestry.

| Disease/PRS | Parameter | EUR |  |  | AFR |  |  |
| --- | --- | --- | --- | --- | --- | --- | --- |
|  |  | AUC (full model) | AUC (covariates only) | AUC difference | AUC (full model) | AUC (covariates only) | AUC difference |
| Type 2 diabetes/<br>Type 2 diabetes | $\rho=0.001$ | 0.5764 | 0.5716 | 0.0048 | 0.7046 | 0.7018 | 0.0028 |
| | $\rho=0.003$ | 0.5769 | | 0.0053 | 0.7067 | | 0.0049 |
| | $\rho=0.01$ | **0.6218** | | 0.0502 | **0.7116** | | 0.0098 |
| | $\rho=0.03$ | 0.6133 | | 0.0417 | 0.709 | | 0.0072 |
| | $\rho=0.1$ | 0.6063 | | 0.0347 | 0.7071 | | 0.0053 |
| | $\rho=0.3$ | 0.6028 | | 0.0312 | 0.706 | | 0.0042 |
| | $\rho=1$ | 0.6015 | | 0.0299 | 0.7056 | | 0.0038 |
| Obesity/<br>BMI | $\rho=0.001$ | 0.6006 | 0.5931 | 0.0075 | 0.5976 | 0.5949 | 0.0027 |
| | $\rho=0.003$ | 0.5991 | | 0.006 | 0.5959 | | 0.001 |
| | $\rho=0.01$ | 0.6453 | | 0.0522 | 0.5981 | | 0.0032 |
| | $\rho=0.03$ | 0.6515 | | 0.0584 | 0.6119 | | 0.017 |
| | $\rho=0.1$ | 0.6539 | | 0.0608 | **0.6119** | | 0.017 |
| | $\rho=0.3$ | **0.6543** | | 0.0612 | 0.611 | | 0.0161 |
| | $\rho=1$ | 0.6529 | | 0.0598 | 0.6099 | | 0.015 |
| Hypertension<br>/<br>Hypertension | $\rho=0.001$ | 0.6979 | 0.6962 | 0.0017 | 0.8418 | 0.8403 | 0.0015 |
| | $\rho=0.003$ | 0.6978 | | 0.0016 | 0.8416 | | 0.0013 |
| | $\rho=0.01$ | 0.6978 | | 0.0016 | 0.8421 | | 0.0018 |
| | $\rho=0.03$ | 0.698 | | 0.0018 | 0.8434 | | 0.0031 |
| | $\rho=0.1$ | 0.7031 | | 0.0069 | 0.8425 | | 0.0022 |
| | $\rho=0.3$ | 0.7096 | | 0.0134 | **0.8435** | | 0.0032 |
| | $\rho=1$ | **0.7143** | | 0.0181 | 0.8434 | | 0.0031 |
| Myocardial<br>Infarction/<br>Myocardial<br>Infarction | $\rho=0.001$ | 0.7686 | 0.7407 | 0.0279 | 0.8355 | 0.8326 | 0.0029 |
| | $\rho=0.003$ | 0.7738 | | 0.0331 | 0.8359 | | 0.0033 |
| | $\rho=0.01$ | **0.7753** | | 0.0346 | 0.8361 | | 0.0035 |
| | $\rho=0.03$ | 0.7721 | | 0.0314 | 0.8369 | | 0.0043 |
| | $\rho=0.1$ | 0.7684 | | 0.0277 | **0.8372** | | 0.0046 |
| | $\rho=0.3$ | 0.7662 | | 0.0255 | 0.8371 | | 0.0045 |
| | $\rho=1$ | 0.7652 | | 0.0245 | 0.8369 | | 0.0043 |
| Coronary atherosclerosis/<br>coronary artery disease | $\rho=0.001$ | 0.7756 | 0.7576 | 0.018 | 0.8425 | 0.8412 | 0.0013 |
| | $\rho=0.003$ | 0.7793 | | 0.0217 | 0.8424 | | 0.0012 |
| | $\rho=0.01$ | **0.7818** | | 0.0242 | 0.8448 | | 0.0036 |
| | $\rho=0.03$ | 0.7794 | | 0.0218 | **0.8448** | | 0.0036 |
| | $\rho=0.1$ | 0.7764 | | 0.0188 | 0.8443 | | 0.0031 |
| | $\rho=0.3$ | 0.7749 | | 0.0173 | 0.8441 | | 0.0029 |
| | $\rho=1$ | 0.7742 | | 0.0166 | 0.8439 | | 0.0027 |
| Renal failure/<br>chronic kidney disease | $\rho=0.001$ | 0.6271 | 0.6271 | 0 | **0.7663** | 0.7657 | 0.0006 |
| | $\rho=0.003$ | 0.6273 | | 0.0002 | 0.766 | | 0.0003 |
| | $\rho=0.01$ | **0.6304** | | 0.0033 | 0.7657 | | 0 |
| | $\rho=0.03$ | 0.6299 | | 0.0028 | 0.766 | | 0.0003 |
| | $\rho=0.1$ | 0.6294 | | 0.0023 | 0.7659 | | 0.0002 |
| | $\rho=0.3$ | 0.6291 | | 0.002 | 0.7658 | | 0.0001 |
| | $\rho=1$ | 0.629 | | 0.0019 | 0.7658 | | 0.0001 |

**Supplemental Table 4:**
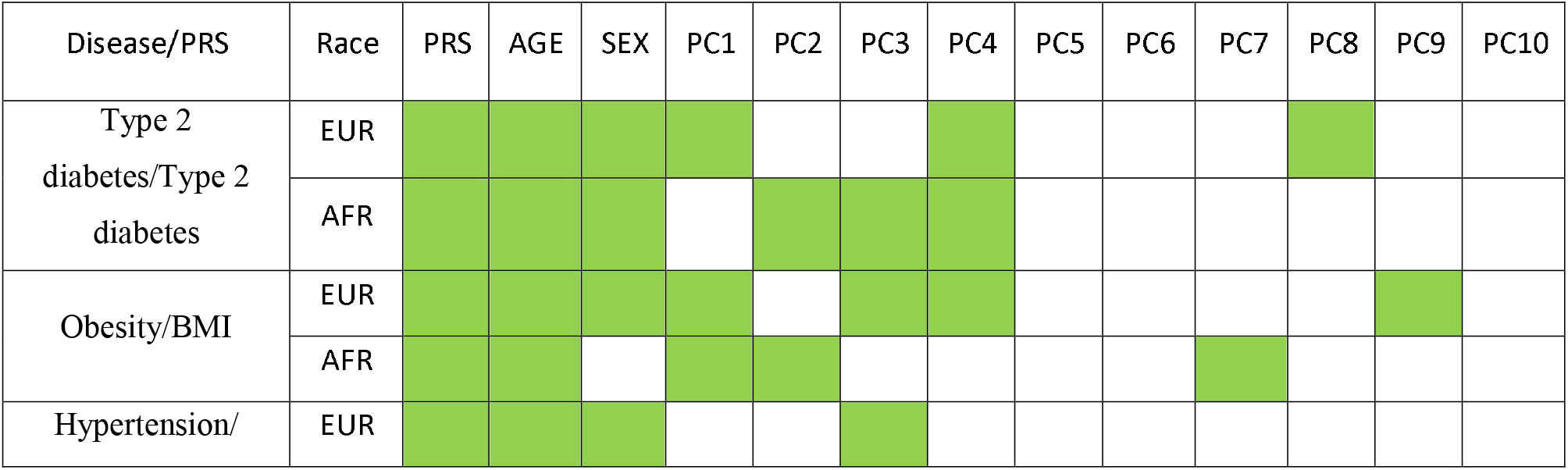
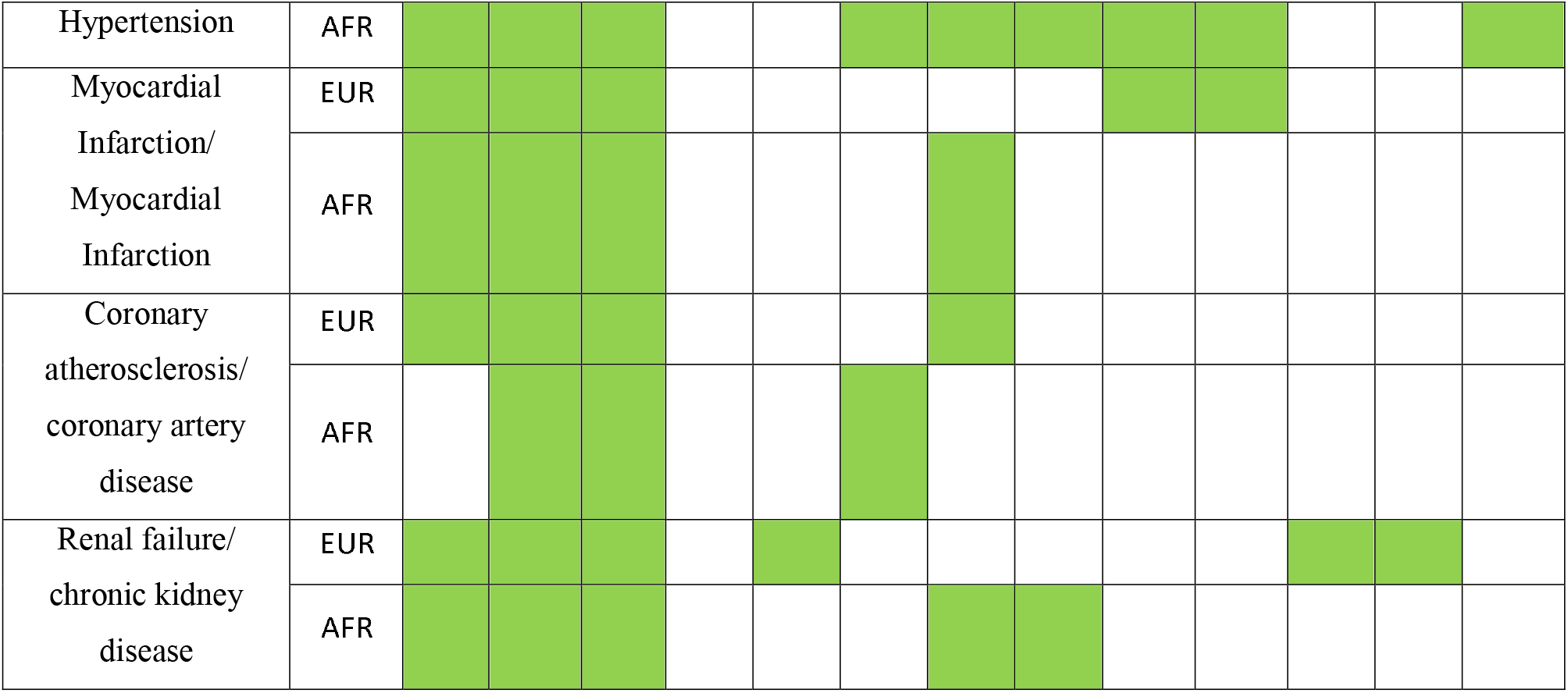
Step-wise regression for model selection. Using the step-wise regression we added each covariate to the null model and colored cells in the table below show covariates selected in best performing regression model. Best model was selected using akaike information criterion (AIC).

**Supplemental Table 5:**
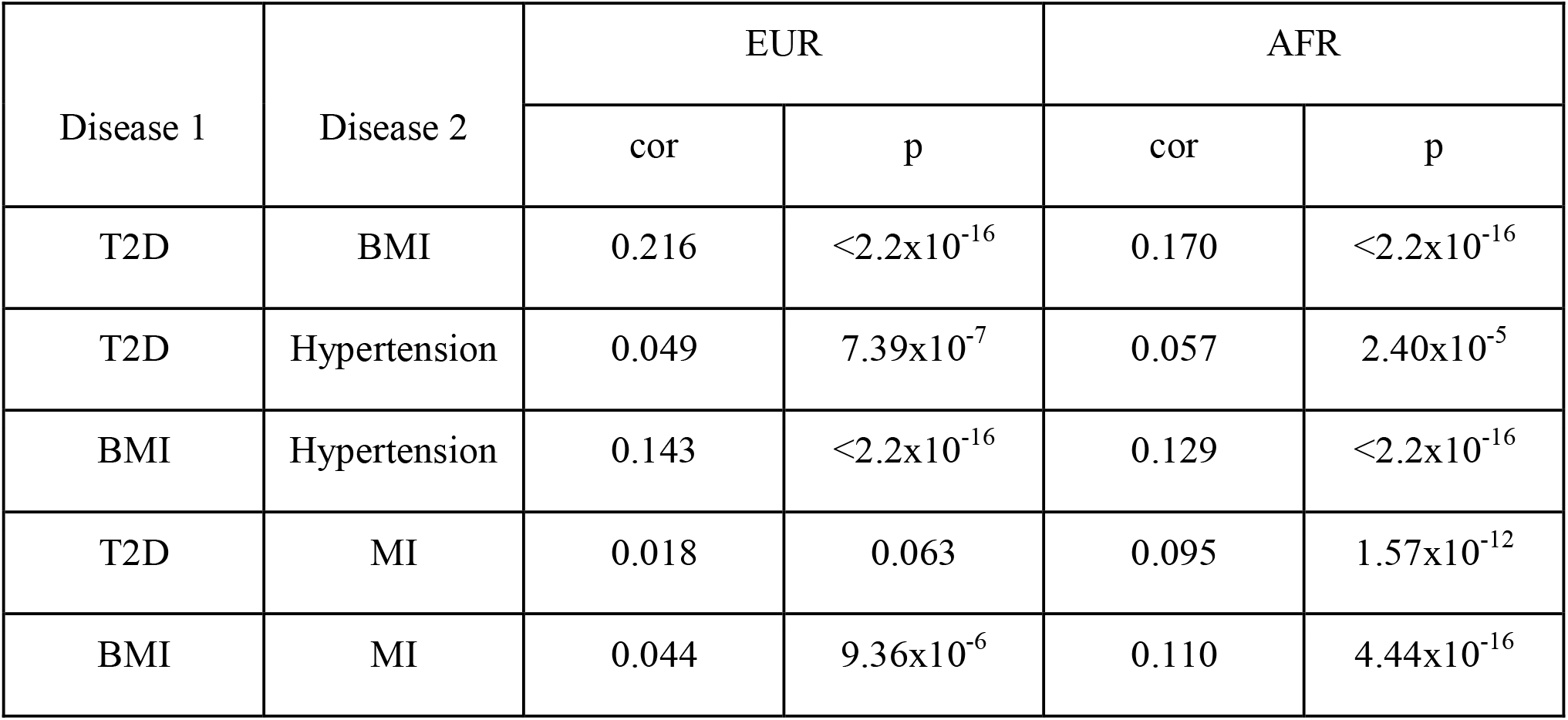
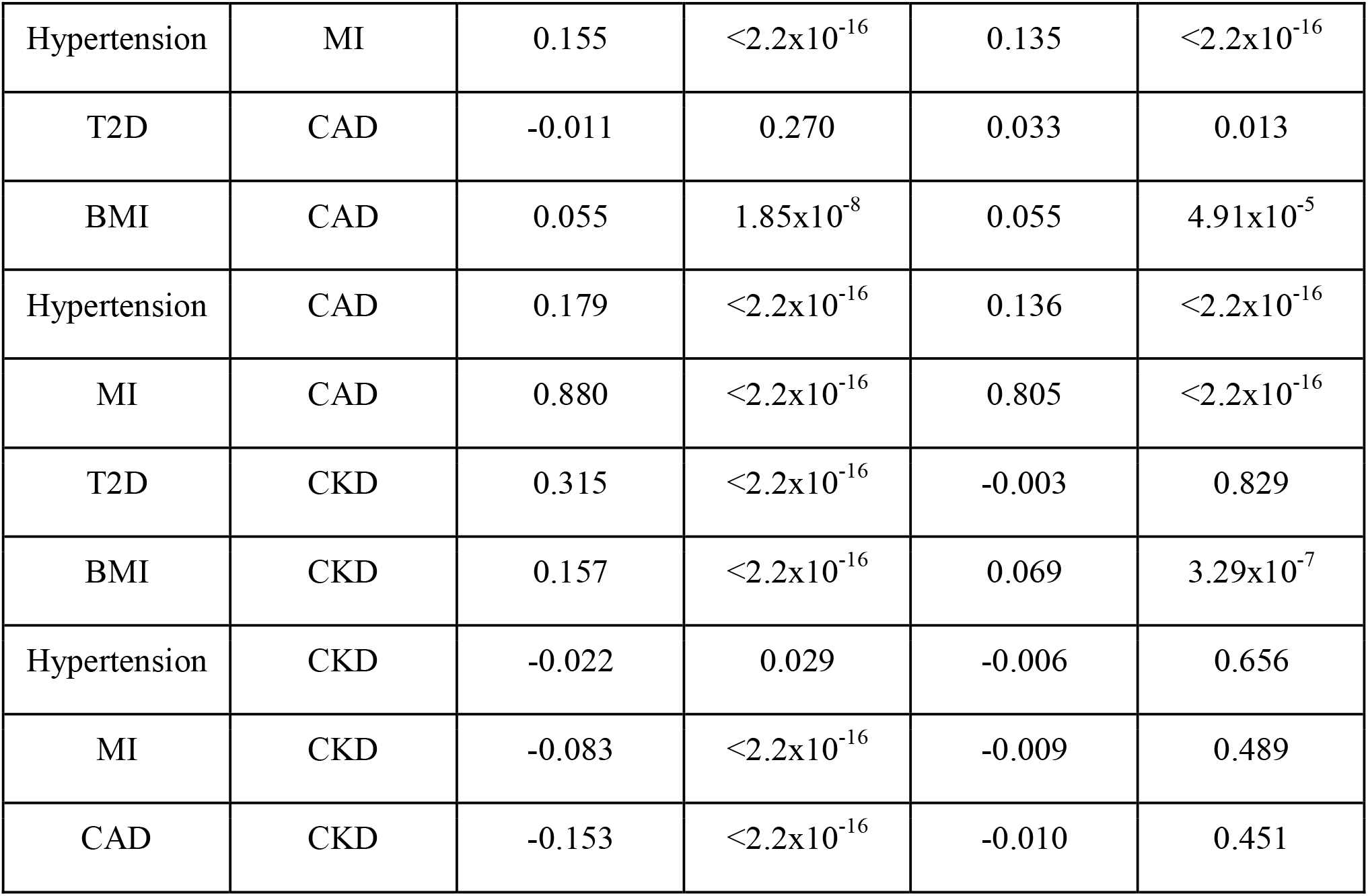
Correlations between PRS.

| Disease 1 | Disease 2 | EUR |  | AFR |  |
| --- | --- | --- | --- | --- | --- |
|  |  | cor | p | cor | p |
| T2D | BMI | 0.216 | $<2.2 \times 10^{-16}$ | 0.170 | $<2.2 \times 10^{-16}$ |
| T2D | Hypertension | 0.049 | $7.39 \times 10^{-7}$ | 0.057 | $2.40 \times 10^{-5}$ |
| BMI | Hypertension | 0.143 | $<2.2 \times 10^{-16}$ | 0.129 | $<2.2 \times 10^{-16}$ |
| T2D | MI | 0.018 | 0.063 | 0.095 | $1.57 \times 10^{-12}$ |
| BMI | MI | 0.044 | $9.36 \times 10^{-6}$ | 0.110 | $4.44 \times 10^{-16}$ |
| Hypertension | MI | 0.155 | $<2.2 \times 10^{-16}$ | 0.135 | $<2.2 \times 10^{-16}$ |
| T2D | CAD | -0.011 | 0.270 | 0.033 | 0.013 |
| BMI | CAD | 0.055 | $1.85 \times 10^{-8}$ | 0.055 | $4.91 \times 10^{-5}$ |
| Hypertension | CAD | 0.179 | $<2.2 \times 10^{-16}$ | 0.136 | $<2.2 \times 10^{-16}$ |
| MI | CAD | 0.880 | $<2.2 \times 10^{-16}$ | 0.805 | $<2.2 \times 10^{-16}$ |
| T2D | CKD | 0.315 | $<2.2 \times 10^{-16}$ | -0.003 | 0.829 |
| BMI | CKD | 0.157 | $<2.2 \times 10^{-16}$ | 0.069 | $3.29 \times 10^{-7}$ |
| Hypertension | CKD | -0.022 | 0.029 | -0.006 | 0.656 |
| MI | CKD | -0.083 | $<2.2 \times 10^{-16}$ | -0.009 | 0.489 |
| CAD | CKD | -0.153 | $<2.2 \times 10^{-16}$ | -0.010 | 0.451 |

Supplemental Table 6: PheWAS of T2D PRS (see excel file)

Supplemental Table 7: PheWAS of BMI PRS (see excel file)

Supplemental Table 8: PheWAS of Hypertension PRS (see excel file)

Supplemental Table 9: PheWAS of MI PRS (see excel file)

Supplemental Table 10: PheWAS of CAD PRS (see excel file)

Supplemental Table 11: PheWAS of CKD PRS (see excel file)

